# A Bioinformatic and Empiric Exploration of Prokaryotic Argonautes as Novel Programmable Endonuclease Systems

**DOI:** 10.1101/2021.03.15.435463

**Authors:** Sejla Salic, Anna Sieber, Samir Barbaria, Anatoly Vasilyev, Elisavet Papadimou, Nevena Milosavljevic, Benjamin Klapholz, Rohan Sivapalan, Trevor Collingwood, Tom Henley, Modassir Choudhry, Tilmann Bürckstümmer

## Abstract

Argonautes are nucleases that can be programmed by short oligonucleotides to cleave complementary sequences. Here, we performed an unbiased bioinformatic search to mine bacterial genomes for prokaryotic Argonautes (pAgos) harboring a PIWI domain. Our search identified 3,033 pAgos in total, of which 1,464 portend to the subgroup of long pAgos with more than 600 amino acids. We purified a subset of 49 pAgos which were found in proximity to helicases and tested their nuclease activity *in vitro*. Ten of these were active towards single-stranded DNA substrates and this activity could be programmed by exogenous guide DNAs or RNAs. Cleavage of double-stranded plasmid DNA was much less readily observed and was fostered by elevated temperatures or exogenous addition of a DNA single-strand binding protein (ET-SSB). The efficiency of pAgo-mediated plasmid cleavage was dependent on the DNA target sequence as well as the surrounding sequence, suggesting that unwinding of the DNA double helix was a limiting factor. Intriguingly, we identified a cluster of pAgos from the Clostridial clade which was active at 37°C and activity was enhanced by exogenous ET-SSB. This suggests that Clostridial pAgos may be particularly suited to catalyze DNA double-strand cleavage and implies that such pAgos may be repurposed as gene editing tools in future.

## INTRODUCTION

The last decade has seen rapid advance in the discovery and characterization of naturally occurring bacterial defense systems that employ DNA or RNA interference mechanisms to protect hosts against invasion by pathogens (Koonin, 2017). The bacterial adaptive immunity system CRISPR/Cas9 has been the most prominent and well-studied of these – due in no small part to a seminal report that elucidated how this system could be recapitulated *in vitro* to function as a programmable nuclease that could be directed to bind to and cleave a desired DNA sequence simply by co-delivering Cas9 nuclease with a guide RNA homologous to the target DNA (Jinek et al., 2012). This elegantly simple system was then readily ported into eukaryotic cells by several groups where it functioned as a highly efficient method for targeted genome modification (Cho et al., 2013; Cong et al., 2013; Jinek et al., 2013; Mali et al., 2013). Such a facile tool for genetic manipulation has since proven to be of immense utility across many fields of application, including human therapeutics and pharmaceutical research, agriculture and industrial biology. Nevertheless, certain limitations of CRISPR/Cas9 such as the prerequisite for a so-called protospacer adjacent motif (PAM) flanking a target site, which limits where Cas9 can bind, together with the large size of most Cas9 proteins, highlights the potential for alternative natural or synthetic systems that circumvent these hurdles to greatly augment the gene editing toolbox.

Argonaute proteins represent a family of nucleases found throughout bacteria, archaea and eukarya (Swarts et al., 2014a). They were first characterized for their role in RNA interference in eukaryotes (Hutvagner and Simard, 2008). In mammalian cells, RNA interference elicited by endogenous miRNAs is believed to contribute to gene regulation and is mediated by a set of mammalian cell-encoded Argonaute genes (Filipowicz et al., 2008). There, Argonautes can load a short single-strand RNA that is typically called short interfering RNA (siRNA) if it is exogenous or microRNA (miRNA) if it is endogenous to the cell. RNA-loaded Argonautes are thus programmed to recognize a complementary RNA in the cell. Upon base pairing, the complementary RNA is cleaved, and its levels are reduced.

In parallel, prokaryotic homologs of Argonaute proteins (pAgos) were predicted to function as key components of a novel defense system against mobile genetic elements (Makarova et al., 2009). Subsequent studies with the prokaryotic pAgo from *Rhodobacter sphaeroides* (RsAgo) showed that it bound both 15-19nt single strand RNA as well as 22-24nt single strand DNA and that RsAgo deficiency led to increased expression of plasmid-encoded genes (Olovnikov et al., 2013). Unlike the CRISPR/Cas9 endonuclease, however, which has two catalytic sites allowing it to cleave both strands of DNA, pAgos have a single site catalytic site and cleave single strand DNA templates. The implication that bacterial Argonautes played a role in eliminating foreign nucleic acids was supported in a subsequent study in which the Argonaute from *Thermus thermophilus* (TtAgo) could cleave a plasmid DNA template *in vitro* (Swarts et al., 2014b). Cleavage was only observed at temperatures >65°C which was attributed to the fact that TtAgo originated from a thermophilic organism and was thus likely to be less active or inactive at lower temperatures. Similar observations were made for additional pAgos, e.g. PfAgo from *Pyrococcus furiosus* (Swarts et al., 2015), and MjAgo from *Methanocaldococcus jannaschii* (Willkomm et al., 2016). More recently, pAgos from several mesophilic bacterial species were characterized in greater detail (Cao et al., 2019; García-Quintans et al., 2019; Hegge et al., 2019; Kuzmenko et al., 2019; Liu et al., 2021). CbAgo from *Clostridium butyricum* and CpAgo from *Clostridium perfringens* can cleave plasmid DNA *in vitro* at 37°C, albeit at limited efficiency. In addition, functional experiments in E.coli suggest that CbAgo can capture single strand guide DNAs from plasmids and phages and can contribute to bacterial host defense (Kuzmenko et al., 2020). This suggests that pAgos can function as programmable nucleases under more mesophilic conditions - at least in bacterial cells.

To extend the universe of pAgos and identify novel nucleases that could be utilized for gene editing, we conducted an unbiased search for novel pAgo sequences. We identified a total of 3,033 pAgo sequences from which we expressed and purified a subset of 49 proteins in E.coli and characterized their ability to cleave single-strand and double-strand DNA *in vitro*. Our approach recovered pAgos that were known to be active but revealed novel pAgos with intriguing functional properties. Moreover, we were able to mitigate the inherent recalcitrance of pAgos towards cleaving double strand DNA at 37°C by inclusion of the single strand DNA binding protein ET-SSB. This raises the possibility of a composite pAgo/auxiliary factor synthetic system as being a solution for gene editing with prokaryotic Argonautes in eukaryotic cells.

## RESULTS

To identify the initial set of candidate pAgos (**Fig. 1A**), we started from all prokaryotic proteins with a PIWI domain annotated in the SMART database (SMART accession number: SM00950). Next, we performed a multiple sequence alignment of the 1,036 PIWI domains found in SMART and used the Jackhmmer tool for a Hidden Markov Model search of the RefSeq Protein Database. This search identified additional pAgos which are not annotated in SMART and expanded our candidate set to 3,033 prokaryotic PIWI-domain-containing proteins (**Supplementary Fig. 1A and Supplementary Table 1**). pAgos have been classified by size as short Agos and long Agos (Ryazansky et al., 2018) and we focused our analysis here on pAgos longer than 600 amino acids (**Supplementary Fig. 1B**). This further narrowed our selection to 1,464 pAgos (**Fig. 1B and Supplementary Table 2**).

**Figure 1.**
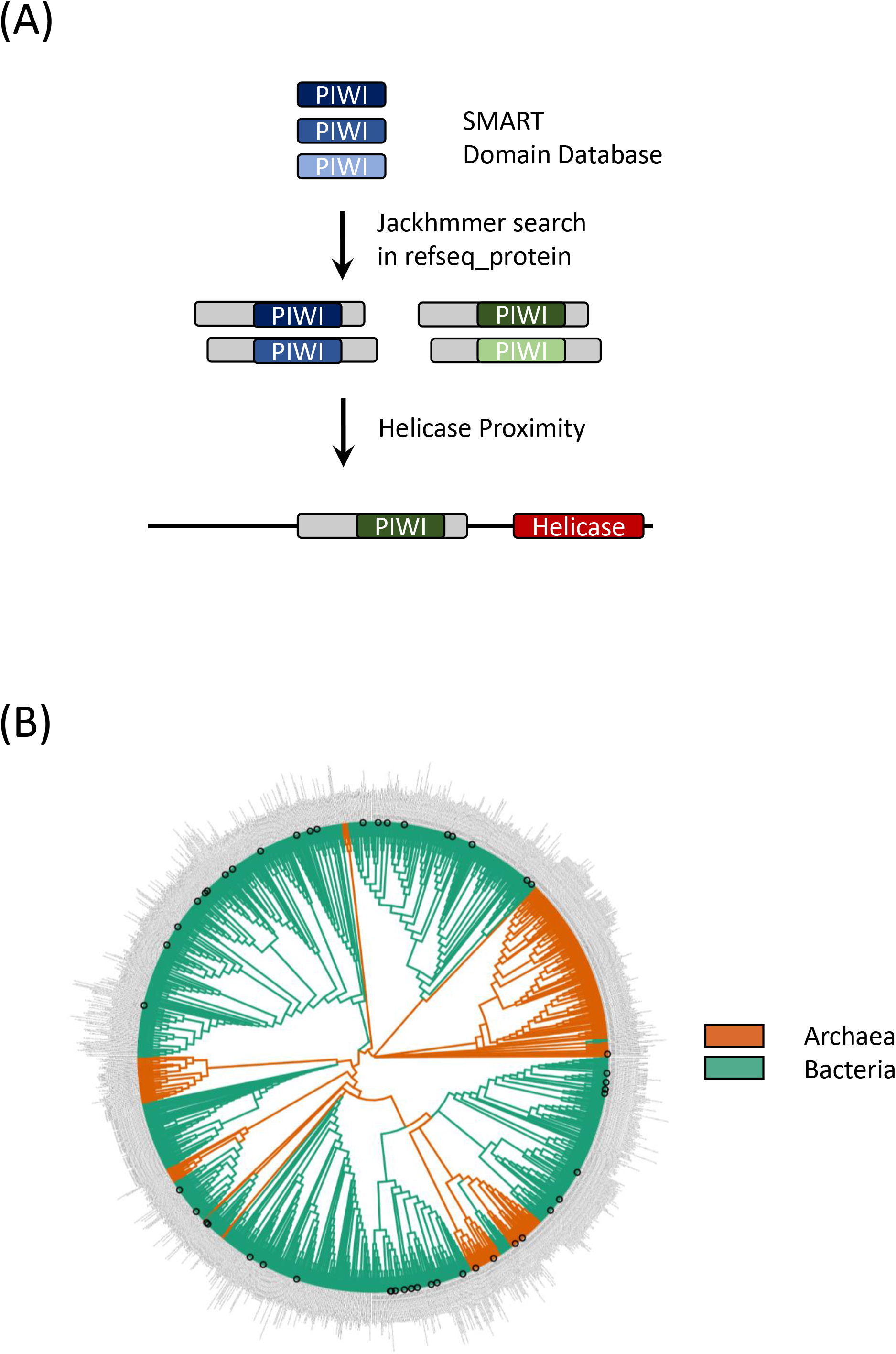
An unbiased bioinformatic search identifies >3,000 prokaryotic Argonautes. (A) PIWI domains of prokaryotic origin were extracted from the SMART database and compared by multiple sequence alignment (MSA). The MSA was used for a Jackhmmer search in the Refseq_protein database, identifying a set of 3,033 prokaryotic proteins bearing PIWI domains. The search was further refined to highlight those pAgos in proximity to helicase domains, thus enriching for pAgos with possible roles in the host defense against DNA. (B) A phylogenetic tree representing the set of 1,464 pAgos and their origin (archaea or bacteria).

A hallmark of prokaryotic operons is the clustering of sequences encoding functionalities required for a given pathway or process (Lee and Sonnhammer, 2003). We reasoned that those pAgos involved in DNA interference or defense against phages or plasmids should either contain or be proximal to functional domains, such as helicases, which facilitate access to the target DNA. Applying those search criteria, we filtered our search to include putative helicase domains within 10 kilobases of the pAgo sequences. Thus, we identified a set of 49 pAgo sequences (**Fig. 2A**) which became the starting point for a detailed experimental characterization.

**Figure 2.**
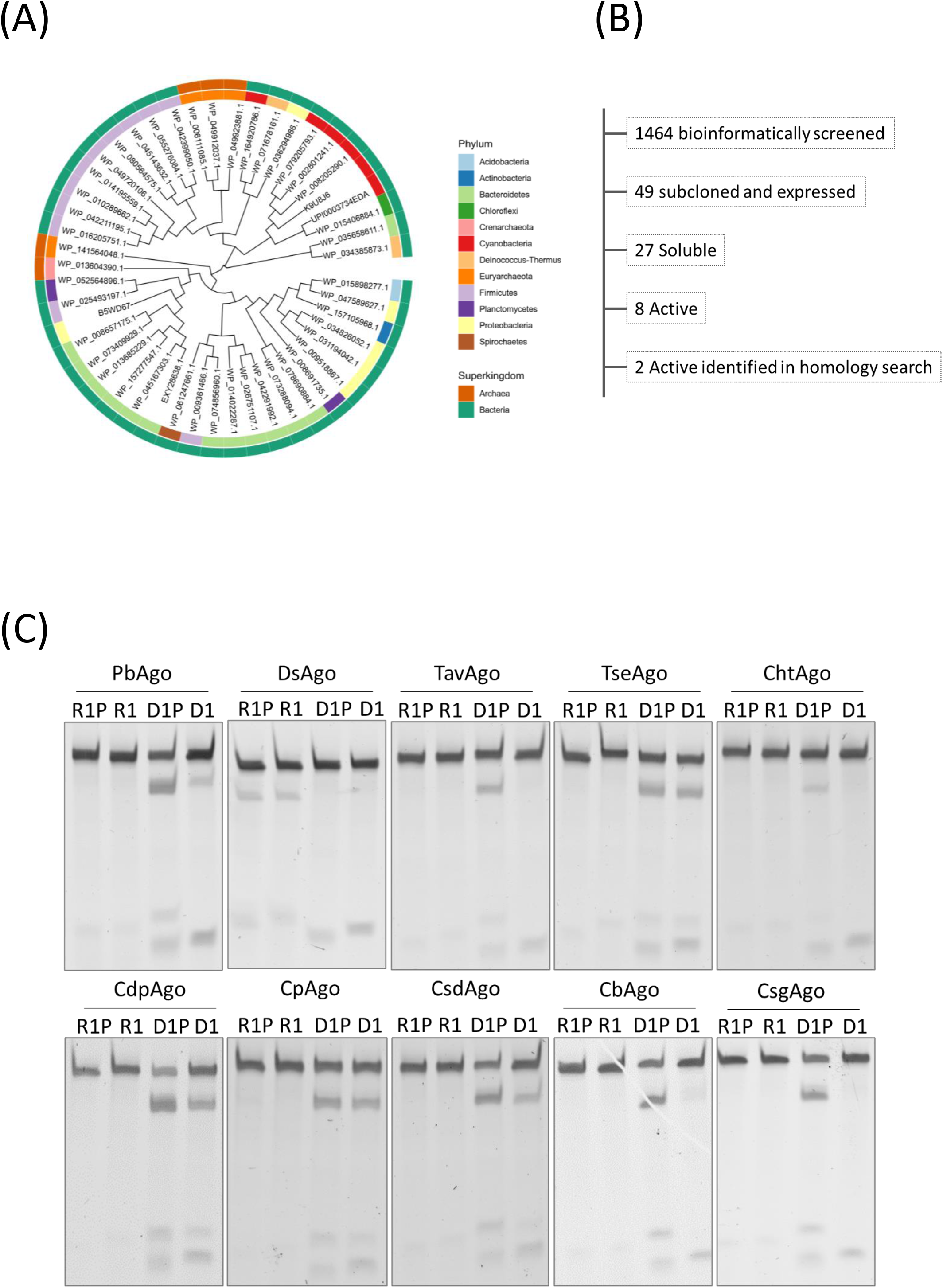
pAgos are potent cutters of single-stranded DNA templates. (A) A phylogenetic tree showing the sequence identifiers and the origin of 49 pAgos used for wet-lab exploration. (B) From all 1,464 candidate pAgo sequences, a set of 49 pAgos were obtained by gene synthesis and cloning. Subsequent expression in E.coli generated a subset of 27 proteins that were expressed and soluble in the conditions used here. A set of 10 pAgos was found to be active on single-stranded DNA (ssDNA) templates. (C) ssDNA cleavage assay of 10 purified pAgos (Table I) incubated with a set of four guides targeting the ssDNA template T1 (**Supplementary Tables 3 and 4**). D1 is unphosphorylated DNA guide; D1P is phosphorylated DNA guide; R1 is unphosphorylated RNA guide, and R1P is phosphorylated RNA guide.

All 49 pAgo-encoding sequences were cloned into a bacterial expression vector and expressed in E.coli. The pAgos were purified initially using single-step NiNTA affinity chromatography and tested for their ability to cleave single strand DNA (ssDNA) *in vitro* at 37°C when paired with phosphorylated guide DNA or RNA. Of the 49 Argonautes screened, 8 emerged that were most active in this primary assay (**Fig. 2B, Table 1 and data not shown**). Interestingly, this set contained three Argonautes from Clostridial species: CdpAgo from *Clostridium disporicum* and CsdAgo from *Clostridium saudiense* which have not previously been studied, and CpAgo which has been shown to be an active programmable nuclease able to cleave plasmid DNA *in vitro* (Cao et al., 2019). Given this seeming enrichment for Clostridial species, we hypothesized that Clostridial Argonautes may be particularly adapted to function as programmable nucleases at mesophilic temperature We therefore performed a homology search for additional proteins closely related to this trio and so identified two more Clostridial Argonautes, namely the previously described CbAgo (Hegge et al., 2019; Kuzmenko et al., 2019) and the uncharacterized CsgAgo from *Clostridium sartagoforme* (**Table 1**).

| Literature name | Species | NCBI accession |
| --- | --- | --- |
| PbAgo | Paenibacillus borealis | WP_042211195.1 |
| DsAgo | Deinococcus sp YIM 77859 | WP_034385873.1 |
| TavAgo | Thermoactinomyces vulgaris | WP_049720106.1 |
| TseAgo | Thermosynechococcus elongatus BP-1 | WP_164920786.1 |
| ChtAgo | Chroococcidiopsis thermalis | uniprot K9U8J6 |
| CdpAgo | Clostridium disporicum | WP_055276084.1 |
| CpAgo | Clostridium perfringens (WAL-14572) | WP_080564575.1 |
| CsdAgo | Clostridium saudiense JCC | WP_042399050.1 |
| CbAgo | Clostridium butyricum | WP_045143632.1 |
| CsgAgo | Clostridium sartagoforme AAU1 | WP_016205751.1 |

The new set of 10 Argonautes was then expressed and purified to >90% purity using a two-step purification protocol (NiNTA affinity chromatography and ion exchange chromatography; **Supplementary Fig. 2**). The pAgos were again tested for their ability to cleave a single-stranded DNA substrate *in vitro* and we compared their activity with 5’-phosphorylated guide DNAs and RNAs to activity seen with their unphosphorylated counterparts. Most of the pAgos showed a preference for guide DNA over guide RNA (**Fig. 2C**), with DsAgo from a *Deinococcus* specie*s* being the only pAgo that displayed selective substrate cleavage with guide RNA. All pAgos we active with phosphorylated guides but only some elicited cleavage with unphosphorylated guides (namely TseAgo from *Thermosynechococcus elongatus*, CdpAgo, CpAgo and CsdAgo).

The PIWI domain of pAgos is considered to contain the catalytic center and displays a conserved DEDX motif (Swarts et al., 2014a). To confirm that the observed cleavage activity was mediated by the PIWI domain we mutated three of the DEDX residues of CpAgo to alanines (D544A, E580A and D730A; **Supplementary Fig. 3A**). Mutant CpAgo was expressed, purified and tested alongside wild-type CpAgo (**Supplementary Fig. 2**) and showed no ssDNA cleavage activity (**Supplementary Fig. 3B**). Similar results were obtained for mutant CdpAgo (harboring the mutations D538A and D608A; **Supplementary Fig. 3C**), confirming that activity as determined here is indeed dependent on the integrity of the PIWI domain. We then assessed the optimal guide DNA length for a subset of the active pAgos (**Supplementary Figure 4A**) and found that an oligonucleotide of 14-15 bases was sufficient to elicit cleavage with only a modest improvement when using longer guides of up to 21 bases. As reported previously for other pAgos (Swarts et al., 2014b), nuclease activity of CpAgo *in vitro* was dependent on the presence of divalent cations and both Mg2+ and Mn2+ could be utilized (**Supplementary Fig. 4B**).

To confirm that substrate cleavage could be triggered in a site-specific manner, we tested four pAgos each with five guide DNAs that targeted different sequences. We observed the expected cleavage pattern with all guides for each of PbAgo (from *Paenibacillus borealis)*, CdpAgo, CpAgo and CsdAgo (**Fig. 3A**). We did note, however, that ssDNA cleavage was almost never complete under these conditions – even upon prolonged incubation at 37°C (data not shown). We hypothesized that secondary structure of the ssDNA target, as predicted by the NUPACK algorithm (**Fig. 3B**), might render the target inaccessible to pAgo cleavage at 37°C. To investigate this, we performed the ssDNA cleavage reaction using CpAgo at temperatures ranging from 25°C to 75°C. We observed greater substrate cleavage efficiency with increasing temperature - achieving almost complete cleavage at 75°C (**Fig. 3C**), in line with the predicted absence of secondary structure at that temperature (**Fig. 3B**). This suggests that secondary structure of a ssDNA target may play a significant role in cleavage by pAgos.

**Figure 3.**
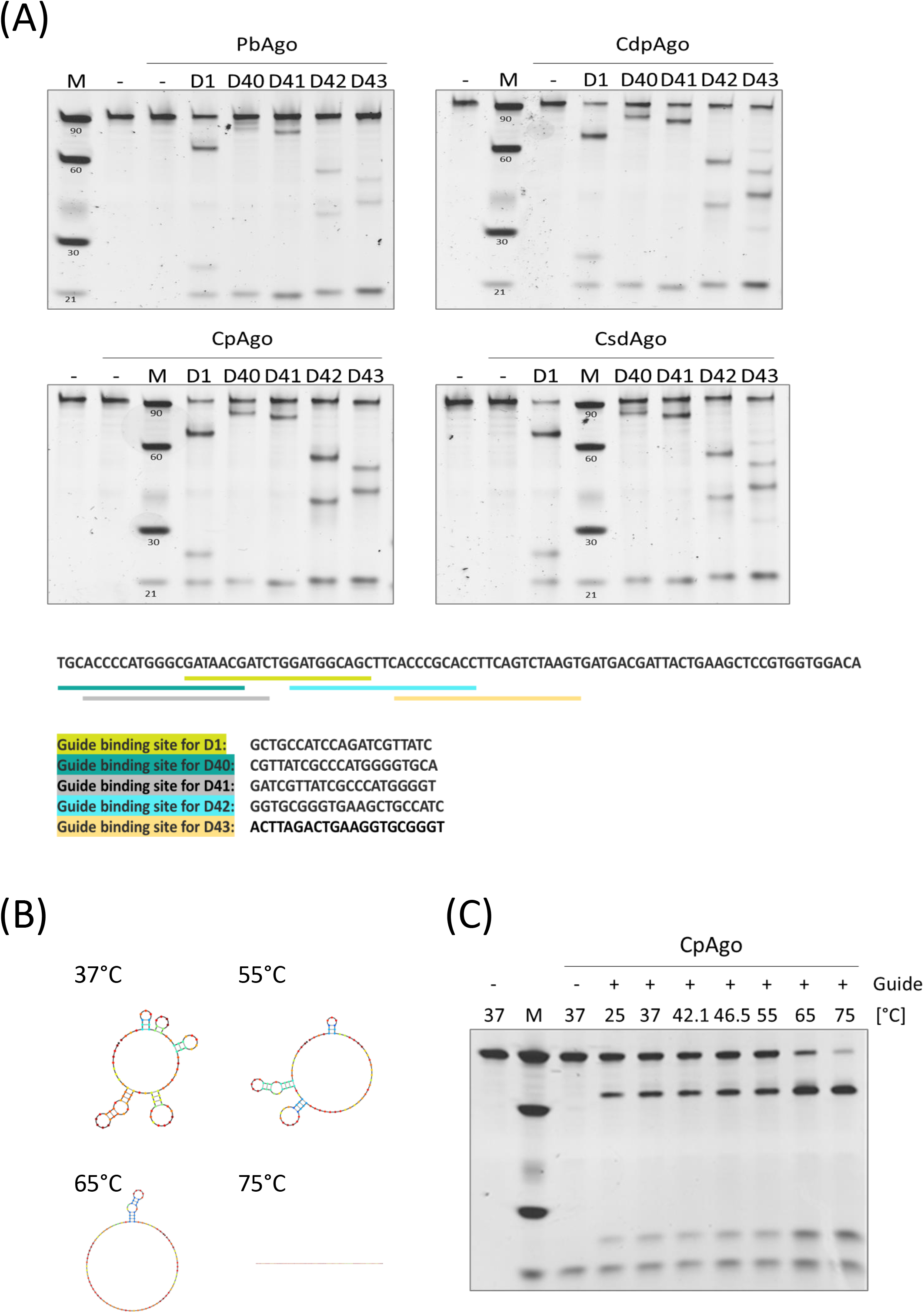
pAgos can be programmed for site-specific cleavage of single-stranded DNA templates. (A) ssDNA cleavage assay of PbAgo, CdpAgo, CpAgo and CsdAgo incubated with a set of five individual guide DNAs (D1, D40, D41, D42 and D42; depicted in color) and a complementary single-stranded DNA (ssDNA) template T2 (**Supplementary Tables 3 and 4**). (B) Secondary structure of the ssDNA template was predicted using the NUPACK tool. (C) ssDNA cleavage assay of CpAgo incubated with guide DNA D1 and a ssDNA template at the temperatures indicated.

A key property of other bacterial defense systems is the ability to elicit targeted cleavage of double strand DNA (dsDNA). In our present study, we first tested CpAgo at a target site in plasmid #79 (**Fig. 4A**). Cleavage of this plasmid was readily detectable when using CpAgo in conjunction with a pair of guide DNAs targeting opposite DNA strands of the target site. In contrast, we found that CpAgo-induced cleavage was not readily detectable at 37°C for a different plasmid #56 (**Fig. 4B**). However, an increase in temperature to 45°C and above led to detectable cleavage of plasmid #56, consistent with the notion that CpAgo may be unable to access the double-stranded DNA of this target at 37°C but is able to bind and cleave individual DNA strands as the duplex separates in response to thermal weakening of strand pairing.

**Figure 4.**
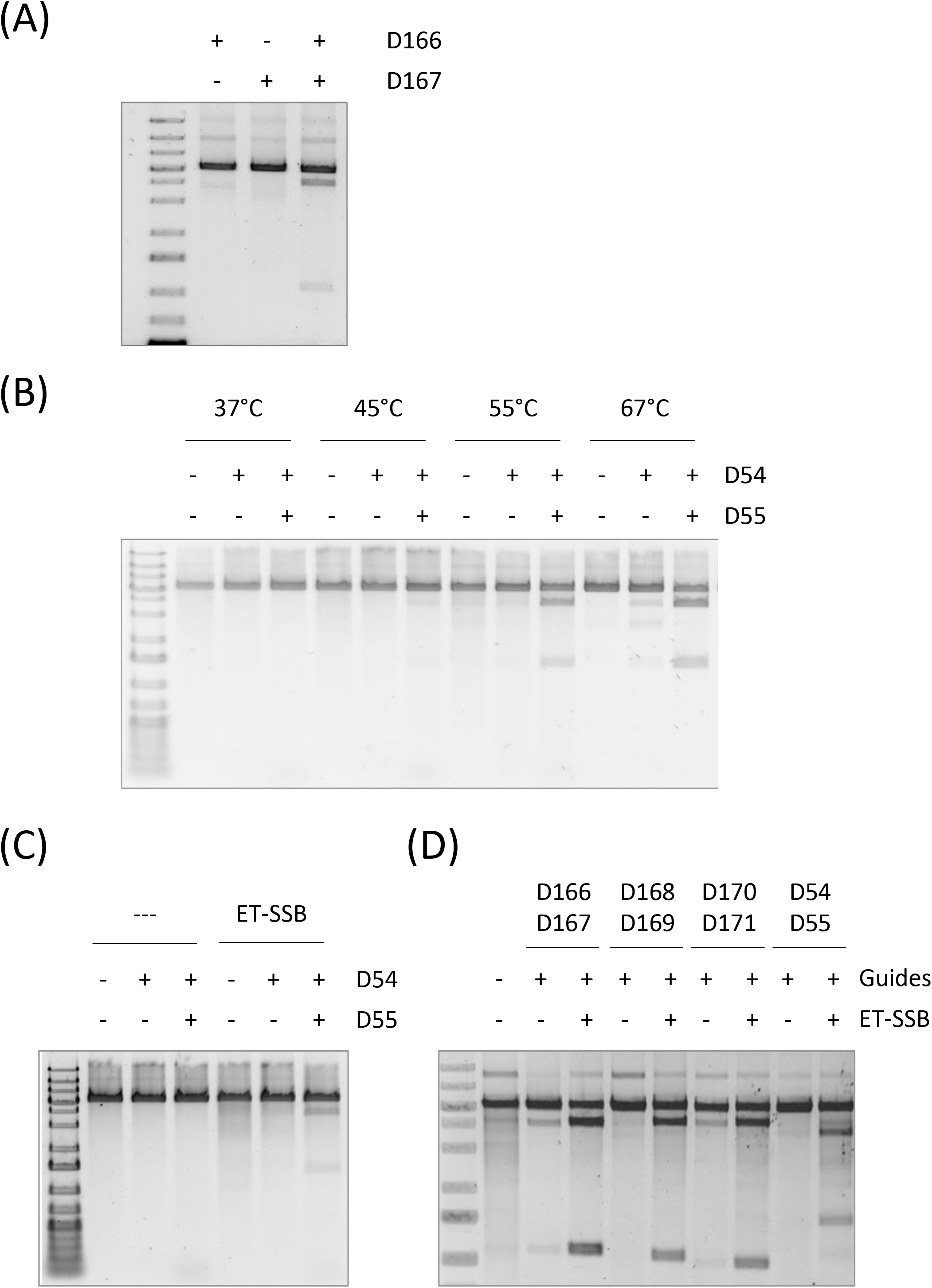
dsDNA cleavage by CpAgo is enhanced by temperature and ET-SSB independently. (A) Cleavage of plasmid #79 at 37°C by CpAgo incubated with a pair of guide DNAs (D166/167; **Supplementary Tables 4 and 5**). (B) Cleavage of plasmid #56 by CpAgo with increasing temperature. CpAgo was incubated with either guide DNA D54 only or a pair of guide DNAs (D54/55). (C) Cleavage of plasmid #56 at 37°C in the presence or absence of ET-SSB. CpAgo was incubated with a pair of guide DNAs (D54/55). (D) Cleavage of plasmid #79 at 37°C with different guide DNA pairs (D166/167; D168/169; D170/171; D54/55) at 37°C in the presence or absence of ET-SSB.

Elevating temperature resolves DNA structural barriers *in vitro* but this approach cannot be applied to cells as it would cause damage and ultimately cell death. We hypothesized that alternative approaches to increasing access to individual DNA strands might provide a solution to facilitating pAgo activity on dsDNA. Indeed, a recent report suggested that addition of single-strand binding (SSB) proteins can facilitate pAgo cleavage *in vitro* (Hunt et al., 2018). SSBs are proteins involved in all aspects of DNA metabolism and stabilize DNA in a single-stranded conformation (Dickey et al., 2013). In line with this, we found that the addition of ET-SSB enabled CpAgo cleavage of the hitherto inaccessible site in plasmid #56 at 37°C (**Fig. 4C**). Moreover, cleavage at this temperature of already accessible sites in plasmid #79 was substantially enhanced (**Fig. 4D**). These results for CpAgo raised the possibility that the inclusion of ET-SSB might be a general strategy for facilitating DNA cleavage by pAgos. We therefore expanded our study to assess the ability of all 10 of the active pAgos from **Fig. 2C** to cleave double-stranded plasmid DNA at 37°C and explored the impact of ET-SSB for each pAgo (**Fig. 5A**). Initially, six of the pAgos, including TavAgo, TseAgo, CdpAgo, CpAgo, CbAgo and CsgAgo showed detectable cleavage of the target site in plasmid #79. Remarkably, four of these six are from Clostridium species – a commensal genus well-adapted to function in the 37°C environment of the mammalian gut (Lopetuso et al., 2013) Cleavage efficiency at 37°C was markedly enhanced by addition of ET-SSB, with two further nucleases, PbAgo and CsdAgo (yet a fifth Clostridial Argonaute) now also showing detectable cleavage, suggesting a general utility for ET-SSB in facilitating pAgo cleavage of dsDNA. Interestingly, cleavage was only moderately further enhanced at 75°C and the elevated temperature did not enable cleavage by any pAgos that were not also active in the presence of ET-SSB at 37°C. Two pAgos (DsAgo and ChtAgo) did not show activity on plasmid DNA under any condition tested here, although both were active on a ssDNA target (**Fig 2C**).

**Figure 5.**
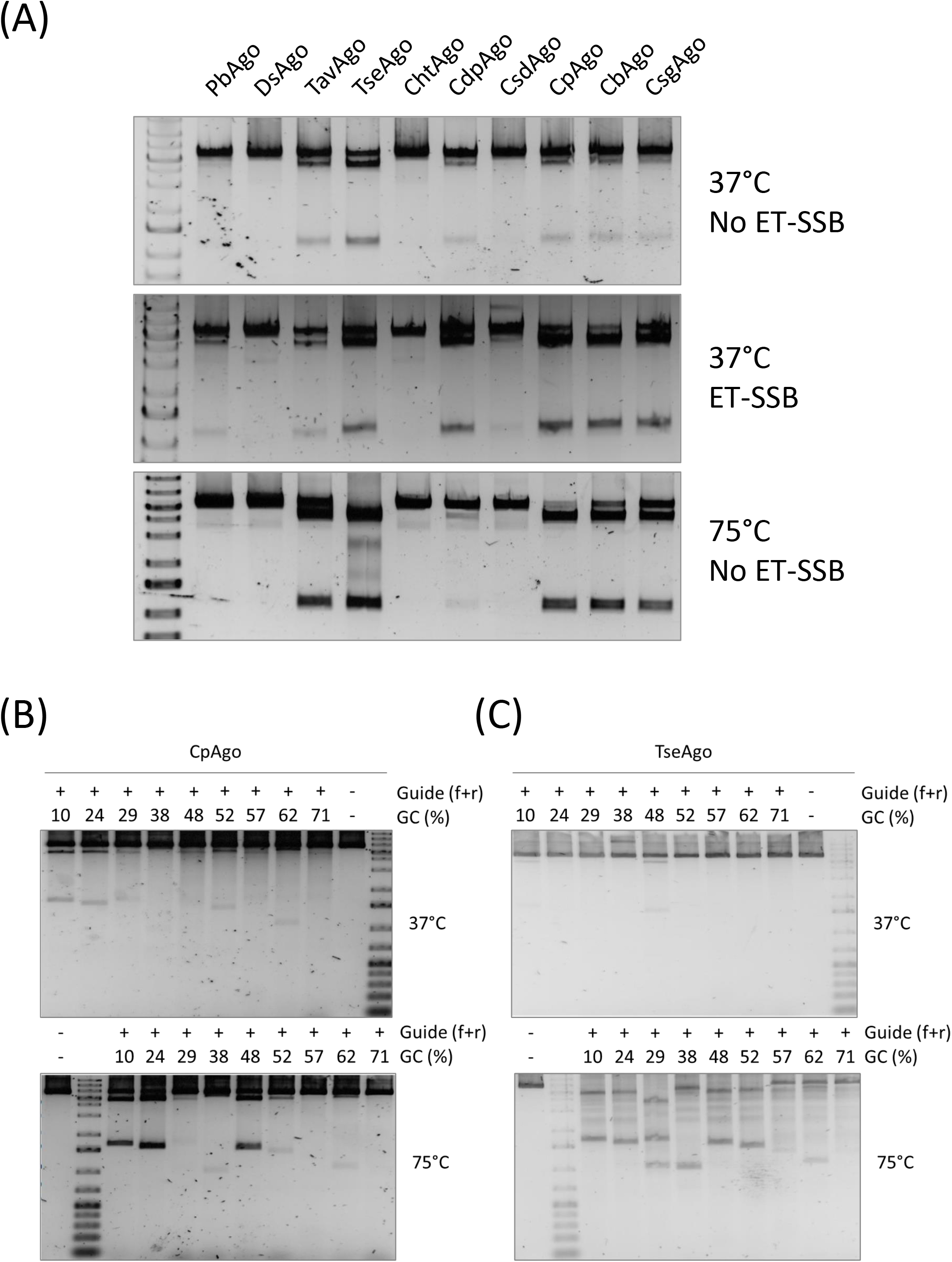
Comparison dsDNA cleavage by ten pAgos. (A) Cleavage of plasmid #79 (guide RNA pair R54/55 for DsAgo and guide DNA pair D166/167 for all other pAgos; **Supplementary Tables 4 and 5**) by 10 pAgos at 37°C in the absence or presence of ET-SSB. Cleavage at 75°C in the absence of ET-SSB is shown for comparison. (B) CpAgo was incubated with plasmid #56 and a set of nine guide DNA pairs harboring increasing GC content in the presence of ET-SSB at 37°C. (C) TseAgo was incubated with plasmid #56 and a set of nine guide DNA pairs harboring increasing GC content in the presence of ET-SSB at 75°C.

The evidence above suggests that improved access of pAgos to the separated strands of DNA is an important determinant of pAgo activity. Previous reports have implicated the GC content of dsDNA target regions as being an important determinant of cleavage efficiency (Cao et al., 2019; Kuzmenko et al., 2019). To further explore the impact of GC content, four of the pAgos (CpAgo, TseAgo, TavAgo, CbAgo) that showed cleavage activity on plasmid DNA at 37°C in the presence of ET-SSB were tested with a set of nine guide DNA pairs ranging in GC content from 10% to 71%. Generally, guide DNA pairs with the lowest GC content (D106/107, 10% GC; D54/55, 24% GC) were most effective at triggering cleavage at 37°C (**Fig. 5B and 5C and Supplementary Figure 5**). However, in the case of CpAgo, cleavage was also seen at 37°C using guide DNA pairs with substantially higher GC content of 29, 52 and 62% (**Fig. 5B)**. Nevertheless, some guide pairs targeting sites with lower GC content were unable to elicit plasmid cleavage by CpAgo at 37°C, indicating that GC content of the target site itself is not the only factor to influence DNA targetability. Cleavage for each pAgo was again enhanced at 75°C but the extent of this temperature effect varied between the pAgos and target site (**Fig. 5B and 5C and Supplementary Figure 5**). Of particular interest however, TseAgo was barely active at 37°C, yet at 75°C it was able to utilize all 8 guide DNA pairs up to 62% GC and thus appears more versatile than the other pAgos tested at that temperature (**Fig. 5C**). However, in the presence of guide DNA, we noted additional cleavage bands for TseAgo that were not observed for the other pAgos, suggesting that TseAgo exhibits promiscuous cleavage activity *in vitro* but only in the presence of guide DNA. Non-specific cleavage by pAgos at 37°C has been reported previously (Swarts et al., 2017) but that was seen in the absence of guide DNA, indicating a different mechanism is likely involved.

The most active pAgo at 37°C, CpAgo, showed plasmid cleavage with certain guide DNA pairs even at relatively high GC content of the target site (62%) yet not for others that had lower GC content **(Fig. 5B)**. This suggested that target site GC content is not the sole determinant of whether a target sequence was cleavable by pAgos and raised the question of whether activity was also impacted by sequence surrounding the target site. To address this question, we created two guide DNA pairs targeting distinct sites in plasmid #56. Guide pair D82/83 (Green) triggered poor CpAgo-mediated cleavage of plasmid #56 at its cognate target site, whereas guide pair D54/55 (Yellow) facilitated high activity at its site (**Fig. 6A and 6B**). In a second plasmid (plasmid #114) the recognition site of D54/55 (the active site in plasmid #56 context) was replaced by the recognition site for D82/83 (inactive in plasmid #56 context), so that this plasmid now bears two recognition sites for the previously inactive D82/83 guides. Upon incubating these plasmids with CpAgo and the respective guide DNA pairs, plasmid #114 could now be cleaved more efficiently using D82/83 (**Fig. 6B**), suggesting that simply shifting the target site for D82/83 to a more permissive location rendered it accessible. Similarly, in a third plasmid (plasmid #115), the recognition sites for D54/55 and D82/83 were swapped with one another – effectively retaining the target sites in the plasmid but changing their surrounding DNA sequence context. Plasmid #115 could no longer be cleaved by D54/55 (the previously active guide DNA pair) once their target site had been moved to the less permissive region previously occupied by the site for D82/83. Both results indicate that pAgo activity is not only dictated by guide DNA sequence, but that the surrounding DNA sequence is also critical – most likely in its impact on DNA target strand separation and allowance of pAgo access.

**Figure 6.**
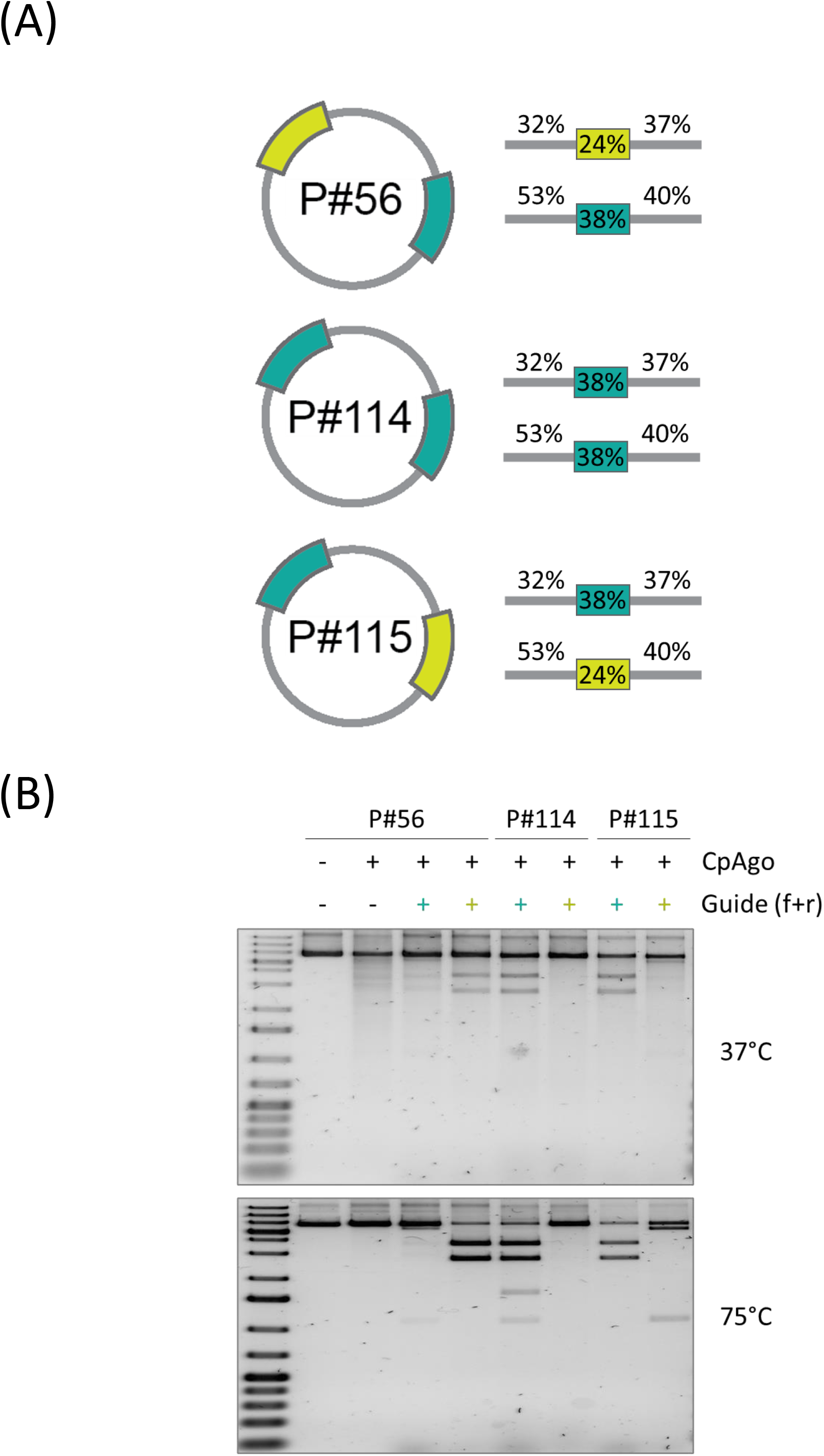
pAgo activity is influenced by sequence surrounding the target site. (A) Schematic representation of the assay design: CpAgo was used to target plasmid #56 using guide DNA pair D54/55 (**Supplementary Table 4**; target site marked in yellow) or guide DNA pair D82/83 (target site marked in green). In plasmids #114 and #115, target sites for guide DNA pairs D54/D55 and D82/83 were swapped as indicated. (B) CpAgo was combined with guide DNA pairs D54/55 and D82/83 as indicated and utilized to cleave plasmids #56, 114 and 115 in the presence of ET-SSB at 37°C or 75°C.

Taken together, our findings show that prokaryotic Argonautes comprise an extended family of programmable nucleases with strong cleavage activity on single-stranded and double-stranded DNA substrates *in vitro* at 37°C. Moreover, cleavage is substantially augmented by exogenous single strand DNA binding proteins that help alleviate barriers to pAgos accessing target DNA. The possibility of now extending the function of this prokaryotic system to the eukaryotic domain offers an intriguing parallel to recent advances in repurposing of a bacterial nuclease-based defense systems to become powerful tools for gene editing.

## DISCUSSION

pAgos are emerging as a class of programmable nucleases that, despite anecdotal published findings, have not been systematically explored. Here, we present a comprehensive study that mined public databases for hitherto unknown or uncharacterized pAgos. The search yielded a large compendium of over 3,000 sequences that we expect will be useful for future functional exploration. In the present study we selected for expression and purification a subset of 49 pAgos of >600 amino acids in length and distributed across the phylogenetic spectrum of bacteria and archaea. Functional characterization identified a remarkable cluster of active pAgos derived from Clostridial bacteria species. While this could represent a bias introduced by our expression host (E.coli) and conditions (37°C), we favor the hypothesis that Clostridial pAgos may indeed be more active at 37°C because Clostridial species are commensal to human physiology and part of the mammalian microbiome. Thus, they have evolved to function at 37°C and may not elicit the same level of antibody response that is found against the *Streptococcus pyogenes* CRISPR/Cas9 system (Charlesworth et al., 2019). Several publications show the potential for Clostridial pAgos to cleavage double-stranded DNA (Cao et al., 2019; García-Quintans et al., 2019; Hegge et al., 2019; Kuzmenko et al., 2019). Very recently, it was shown that CbAgo can function in the defense against plasmids or phages (Kuzmenko et al., 2020). Importantly, this activity was portable to E.coli, which represents a key prerequisite for the re-purposing of pAgos as gene editing tools.

### pAgos show distinct requirements for guide structure

One of the notable distinctions between prokaryotic Argonautes and those from eukaryotes is the general preference for the latter to use short single stranded RNA molecules to guide target recognition. In contrast, most pAgos studied to date have been found to prefer single stranded DNA guides. In the present study, we found that nine of the ten pAgos examined require single strand DNA guides to efficiently cleave their DNA targets, with only DsAgo demonstrating preference for guide RNA (**Table 1**). Of note, too, was the selective importance of 5’ phosphorylation of the guides. Six of the pAgos (PbAgo, DsAgo, TseAgo, CdpAgo, CpAgo and CsdAgo) exhibited a mild to moderate preference for the phosphorylated form of guide, while four (TavAgo, ChtAgo, CbAgo and CsgAgo) showed an absolute requirement for 5’ phosphorylation. These results are consistent with previous studies on CbAgo and CpAgo (Cao et al., 2019; García-Quintans et al., 2019; Hegge et al., 2019), though another report suggested CbAgo had minimal preference for 5’ phosphorylation (Kuzmenko et al., 2019). Despite the seemingly rigid constraints regarding phosphorylation status in some instances, there was however greater latitude for guide length which showed similar efficiency over a range of 14-21 nucleotides, as also seen by others (Cao et al., 2019; Hegge et al., 2019; Kuzmenko et al., 2019), although one study showed cleavage of a 45-nucleotide ssDNA template by CbAgo with a guide as short as 7 nucleotides (García-Quintans et al., 2019). Future structural analyses of this cohort of guide-bound pAgos and others will help elucidate the subtle variations in molecular mechanisms by which guide structure drives pAgo activation.

### DNA target structure is a key determinant of pAgo function

pAgos are clearly very effective at cleaving single strand DNA templates *in vitro*, while double stranded DNA represents a greater challenge. Nevertheless, we have shown here that even ssDNA target structure can present a barrier to pAgo cleavage - likely through the formation of secondary structures that block access by the pAgo/guide complex. Indeed, a comparison of NUPACK structure predictions of single strand DNA templates used by us and by others indicated a correlation between the completeness of template cleavage and the expected presence of secondary structure at a given temperature (data not shown). It seems, however, unlikely that secondary structure of a single strand of DNA would be a critical factor in the context of a potential genomic DNA target. Of greater importance will be the ability of a pAgo complex to access the genomic DNA duplex. Studies to date have use supercoiled plasmid DNA as model template for dsDNA cleavage (Cao et al., 2019; Swarts et al., 2014b). While modest cleavage of plasmid could be achieved at 37°C on a template with particularly low GC content around the target site, cleavage of higher GC content targets required pre-incubating the dsDNA at temperatures substantially higher than 37°C. This was consistent with the notion that higher temperatures drive increased strand separation of the DNA such that pAgos can then effectively create a “double strand break” in dsDNA by using two separate guide/pAgo complexes targeted to and cleaving opposing strands of the DNA. This is in contrast to CRISPR/Cas9 where one nuclease molecule uses its two catalytic domains to create a double strand break. In the present study, we too initially observed relatively low levels of plasmid cleavage at 37°C by the previously characterized CpAgo and CbAgo, but we have extended such findings to now include the four novel nucleases: TavAgo, TseAgo, CdpAgo and CsgAgo **(Fig. 5A**). Thermal denaturation of the target DNA at 75°C facilitated cleavage for five of the pAgos – with the exception being CdpAgo (**Fig. 5B and 5C)**. The surprisingly versatile targeting profile of TseAgo at 75°C indicates that there are fundamental differences between pAgos that can result in quite diverse properties. The harnessing and optimizing of a nuclease like TseAgo might therefore offer an intriguing route to a pAgo-based gene editing system that has fewer limitations arising from target DNA structure.

The set of ten pAgos tested here all showed high programmable nuclease activity towards single-stranded DNA substrates and all guide DNAs (and guide RNAs for DsAgo) tested conferred activity. In contrast, only certain guides could be utilized for programmed cleavage of plasmid DNA. This suggests that it is not the inherent inability of the pAgos to bind certain guides, or to mediate hydrolysis of the phosphodiester bond, that determines whether a pAgo will cleave its target. Rather, it implies that certain regions of a plasmid may be less accessible for cleavage and it has been proposed that the GC content of a target site is an important determinant of pAgo activity (Cao et al., 2019; Hegge et al., 2019; Kuzmenko et al., 2019). However, we showed that while, in general, lower %GC sites proved easier to target, GC content of the immediate target was not a sure predictor of likely success (**Fig. 6**) and that the sequence context of the target site is also critical.

## ET-SSB as a strategy to enhance pAgo function at 37°C

A requirement for low GC content of the target site or the use of temperatures well above 37°C presents a significant limitation to adapting pAgos to cleave dsDNA in the context of a eukaryotic genome. To mitigate the challenge of accessing DNA inside a cell, we made use of the notion that pAgos often colocalize with other DNA-opening functionalities in the same or neighboring operons and that auxiliary factors such as helicases or single strand DNA binding proteins might be employed naturally to aid DNA strand separation at mesophilic temperatures (Hunt et al., 2018; Swarts et al., 2014a). To this end, we investigated several single strand DNA binding proteins (SSBs) and helicases for their ability to facilitate pAgo-mediated cleavage of plasmid DNA *in vitro* and identified ET-SSB as the most favored candidate (**Fig. 4C** and data not shown). In some cases, the robust positive effect of ET-SSB on plasmid cleavage approached that of elevating the cleavage temperature to 75°C (**Fig. 5A)**. From the perspective of using ET-SSB to facilitate cleavage of higher GC target sites at 37°C, the activity of CpAgo in particular was most encouraging (**Fig. 5B**). Future investigation may refine a subset of pAgos that, together with ET-SSB or other auxiliary factors, can be repurposed for genome manipulation in a broad array of cell types.

A core attribute of an efficient and useful gene editing system is a precise DNA domain binding element. As such, this Argonaute-based programable endonuclease system may serve as the basis for a modular system of site-specific double strand DNA binding from which advanced genome editing applications – in particular, base editing and transposon-coupled integration - can follow. The essential element as described in this paper is the elucidation of a programmable DNA binding motif and augmentation of its function. The next step is to extend this exploration into establishing DNA binding in the eukaryotic genome. In summary, our studies suggest a path forward involving the combination of pAgos with auxiliary factors such as SSBs or DNA unwinding proteins – possibly as fusions with pAgos - by which pAgos might function effectively on dsDNA without the need for temperatures beyond the physiological boundaries of mammalian cells. This may prove a vital step in advancing this system into eukaryotes as a valuable new gene editing tool.

## MATERIALS AND METHODS

### Bioinformatic Identification of PIWI domain containing proteins

First, all the available prokaryotic PIWI domain sequences were downloaded from the SMART database (Letunic et al., 2020). This yielded 1,036 sequences. We then sought to identify all the proteins potentially containing this domain. The full set of protein sequences (non-redundant) found in Bacteria and Archaea was downloaded from the RefSeq database. A multiple sequence alignment of the 1,036 PIWI domain sequences was constructed using *Clustal Omega* (Madeira et al., 2019) with default parameters and used as an input to *jackhmmer* (v. 3.3.2; http://hmmer.org/) tool to search for this domain in RefSeq proteins. Separate runs for Bacterial and Archeal sequences were performed, both with default parameters except *--tblout* (tabular output) and *--domE 0*.*001* (significance filtering) and in the case of bacterial proteins, *-N 20* (algorithm iterations). *jackhmmer* search was interrupted as soon as the rate of new hits being found dropped significantly. This search yielded 2,722 sequences from RefSeq, with certain overlap to the original set of 1,036 PIWI-containing proteins from the SMART database. The union of these two sets yielded 3,033 sequences with unique NCBI accession numbers. Finally, this set was narrowed down to 1,464 candidate proteins by only selecting sequences in the size range of 600 to 1100 amino acids.

### Phylogenetic tree visualizations

All the phylogenetic tree visualizations in this publication were created as follows: a multiple sequence alignment was created using *AlignSeqs* function from *DECIPHER* (Wright, 2016) package (v. 2.18.1) in R (v. 4.0.3; http://www.R-project.org/.) with default parameters and exported into *phylip* format. The phylogenetic tree was then built using *FastTree* (v. 2.1.11) with default parameters and visualized using *ggtree* package (v. 2.4.1) from R.

### Protein expression and purification

Synthetic genes each encoding one of the listed 49 prokaryotic Argonautes were ordered from Twist Bioscience and inserted into a pETM30 expression vector in frame with the N-terminal His-GST tags. The subcloned AGO plasmids were transformed into *Escherichia coli* BL21 (DE3) (New England Biolabs) according to manufacturer’s instructions. Strains were cultivated in LB medium (Carl Roth) containing 50 μg/ml Kanamycin (Carl Roth) in a bacterial shaking incubator at 37°C and 150 rpm. After overnight incubation, the preculture was used to inoculate expression cultures with a starting OD600 of 0.05. The cultures were grown at 37°C and 150 rpm until OD600 of 0.8 was reached. Protein expression was induced by adding 1 mM of isopropyl-β-D-thiogalactoside (IPTG) (Sigma Aldrich). Expression was continued in a bacterial shaker at 30°C and 150 rpm for 6h. Cells were harvested by centrifugation at 5000 x *g* for 10 min at 4°C. The pellet was frozen and stored at −80°C. The frozen cells were thawed at 4°C and resuspended in 25 mL buffer I (50 mM Tris/HCl pH 7.5, 0.5 M Sodium chloride, 5% Glycerol) supplemented with 1 mM Phenylmethylsulfonyl fluoride (Carl Roth) and 5 mM β-Mercaptoethanol (Sigma Aldrich). The resuspended cells were disrupted by sonication with a Branson Digital Sonifier (Model 102C, 3 mm tip). Sonication: Step 1: 25% amplitude; 5 sec ON, 2 sec OFF for 2 min; repeat twice; pause for 3 min after each cycle; Step 2: 35% amplitude; 5 sec ON, 2 sec OFF for 30 sec. The lysed pellet is kept on ice during sonication. The lysate was centrifuged for 15 min at 15000 x *g* at 4°C, after which the soluble fraction was used for His-Tag affinity chromatography purification. The cleared lysate was incubated with Ni-NTA agarose on a rotary wheel (30 min at 4°C). After centrifugation (50 x *g* for 5 min) the Ni-NTA agarose beads were transferred to an empty Bio-Spin Chromatography column (Biorad). The column was extensively washed with 20 CV (column volume) of buffer I supplemented with 5 mM β-Mercaptoethanol. The tagged protein was gradually eluted with buffer I supplemented with 5 mM β-Mercaptoethanol and increasing concentrations of Imidazole (25 mM, 50 mM, 75 mM, 125 mM, 3×250 mM. Fractions 3-7 were pooled and diluted 1:5 in buffer III (50 mM Tris/HCl pH 7.5, 150 mM NaCl, 5% Glycerol, 5 mM β - Mercaptoethanol). The His-GST tags were cleaved by incubating the pooled fractions with TEV protease (New England Biolabs) at 4°C for 16h. TEV protease and cleaved tags were removed by performing a reverse NiNTA batch purification. The supernatant was purified over a cation exchange column (GE Healthcare) using an ÄKTA FPLC system. Proteins were dialyzed overnight against buffer IV at 4°C. Aliquots were frozen in liquid nitrogen and stored at −80°C. Mutant CpAgo (D544A, E580A, D730A) and mutant CdpAgo (D538A and D608A) cDNAs were synthesized by Twist Bioscience and subcloned into the pETM30 expression vector by Gibson assembly (New England Biolabs).

### DNA cleavage assays on single-stranded DNA templates

Single-stranded DNA activity assays were performed at equimolar amounts of pAgo protein, guide and target DNA in 1x CutSmart buffer (New England Biolabs). The final concentration of protein, guide and DNA template in a total reaction volume of 15 µl is 250 nM. Proteins were prediluted to a concentration of 750 nM in 1x CutSmart buffer (New England Biolabs) and preloaded with guide DNA or guide RNA in a 1:1 ratio at 37°C for 15 min. The, single-stranded DNA template is added and incubated at 37°C or as indicated for 1h. To inactivate ssDNA cleavage reactions, samples were incubated with TBE urea sample buffer (Biorad) in a 1:1 ratio at 95°C for 10 min. ssDNA cleavage products were resolved on 15% TBE Urea gels (Invitrogen). As a marker, an in-house prepared ssDNA marker consisting of 21, 30, 60 and 90 nt oligos was used. Gels were stained for 10 min with SYBR gold Nucleic Acid Gel Stain (Invitrogen) and visualized using a UVsolo TS Imaging System (Analytik Jena). Sequence of the ssDNA templates T1 and T2 is shown in **Supplementary Table 3**. Sequence of the guide DNA and guide RNA is shown in **Supplementary Table 4**.

### DNA cleavage assays on double-stranded DNA templates (plasmids)

The preloading with forward or reverse guide DNA/RNA was carried out in separate reactions. If not indicated otherwise, reactions were performed in 1x CutSmart buffer (New England Biolabs). Proteins were prediluted to a final concentration of 750 nM in 1x CutSmart buffer (New England Biolabs) and incubated with the respective guide at 37°C for 15 min in a 1:1 ratio. Next, half reactions of preloaded protein were combined, and ET-SSB (New England Biolabs) as well as 200 ng of target plasmid DNA was added and incubated at 37°C (or other temperature) for 1h. Subsequently, the reaction volume was adjusted to 28 µl with 1x CutSmart buffer and 1 µl of the indicated restriction enzyme was added. The reaction was incubated at 37°C for 1h. dsDNA cleavage assay reactions were inactivated with Proteinase K solution (1µl; 20 μg/reaction) (Qiagen) for 20 min at room temperature. Samples were mixed with 6x loading dye (New England Biolabs) before they were resolved on a 1% agarose gel. As a marker, a 1kb Generuler Marker was used (Thermo). Gels were stained for 10 min with SYBR gold Nucleic Acid Gel Stain (Invitrogen) and visualized using a UVsolo TS Imaging System (Analytik Jena). Sequence of the guide DNA and guide RNA is shown in **Supplementary Table 4**. Plasmid templates are described in **Supplementary Table 5**.

## SUPPLEMENTARY FIGURE LEGENDS

**Supplementary Figure 1.**
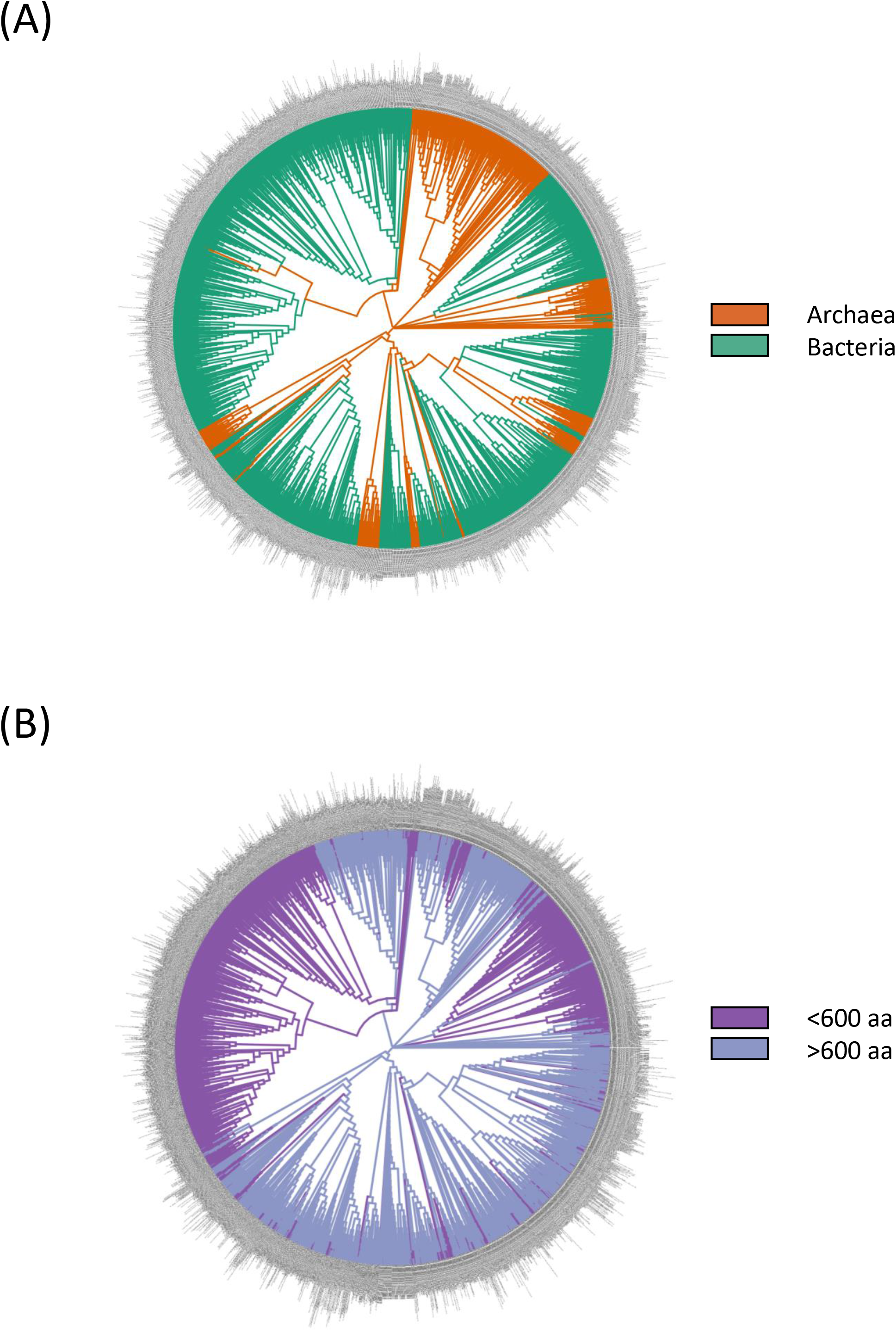
Phylogenetic relationships of prokaryotic Argonautes. A set of 3,033 prokaryotic Argonautes was obtained as described in Materials & Methods. In brief, PIWI domain-containing proteins were obtained from the SMART domain database. Additional such proteins were obtained by mining the RefSeq protein database using the jackhmmer tool. The phylogenetic tree was colored by (A) origin (orange for archaea and green for bacteria) or by (B) length of the prokaryotic Argonautes obtained (<600 amino acids colored in purple, >600 amino acids colored in blue).

**Supplementary Figure 2.**
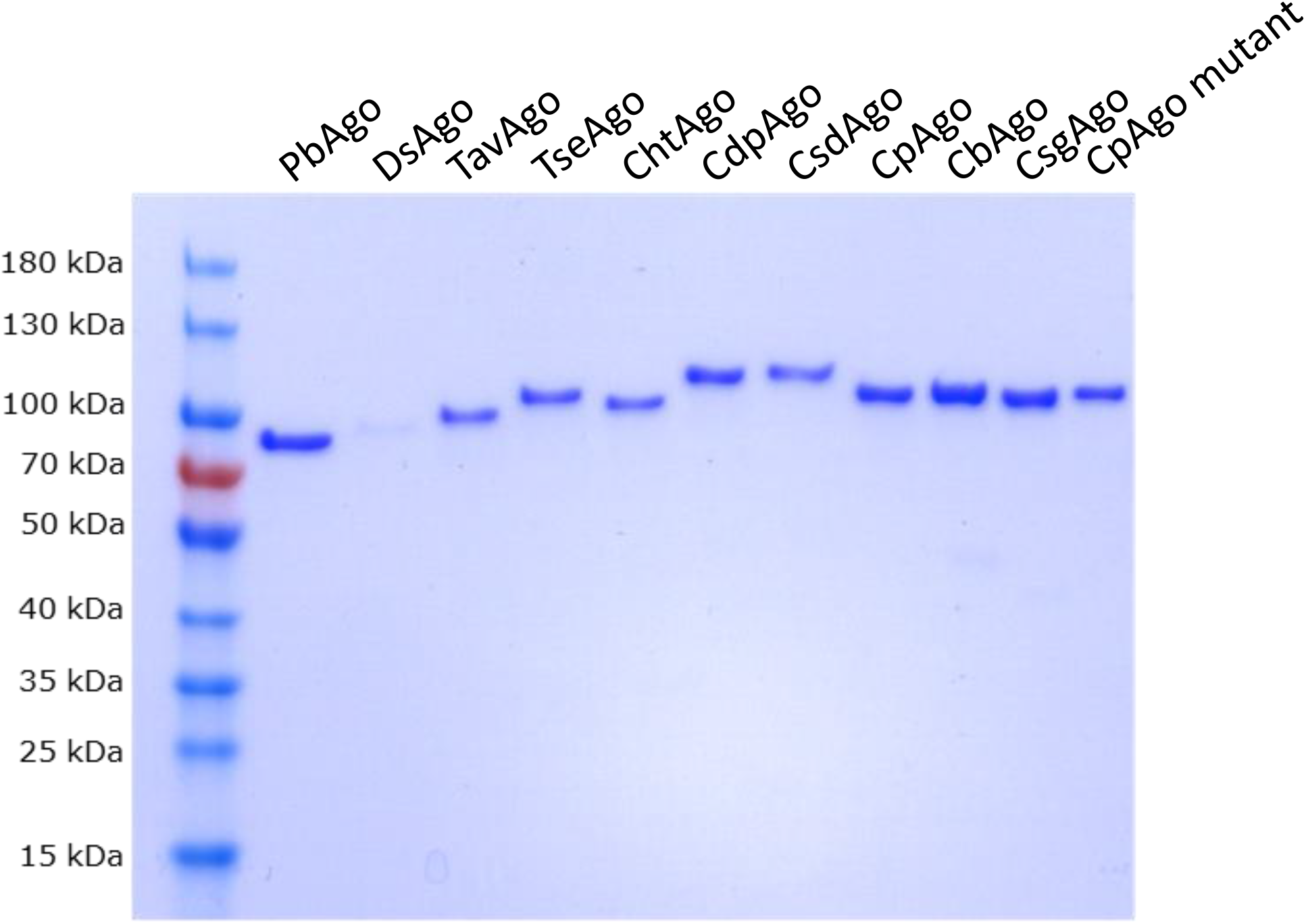
Recombinant prokaryotic Argonaute were purified to homogeneity. Recombinant pAgos were expressed and purified as described in Materials & Methods. In brief, pAgos were expressed as GST fusion proteins in BL21 bacteria from a pET-based vector. Lysates were purified by NiNTA chromatography. Next, the GST fusions were cleaved using TEV and the GST was removed by another NiNTA chromatography set. In a final step, the protein was polished by cation exchange chromatography.

**Supplementary Figure 3.**
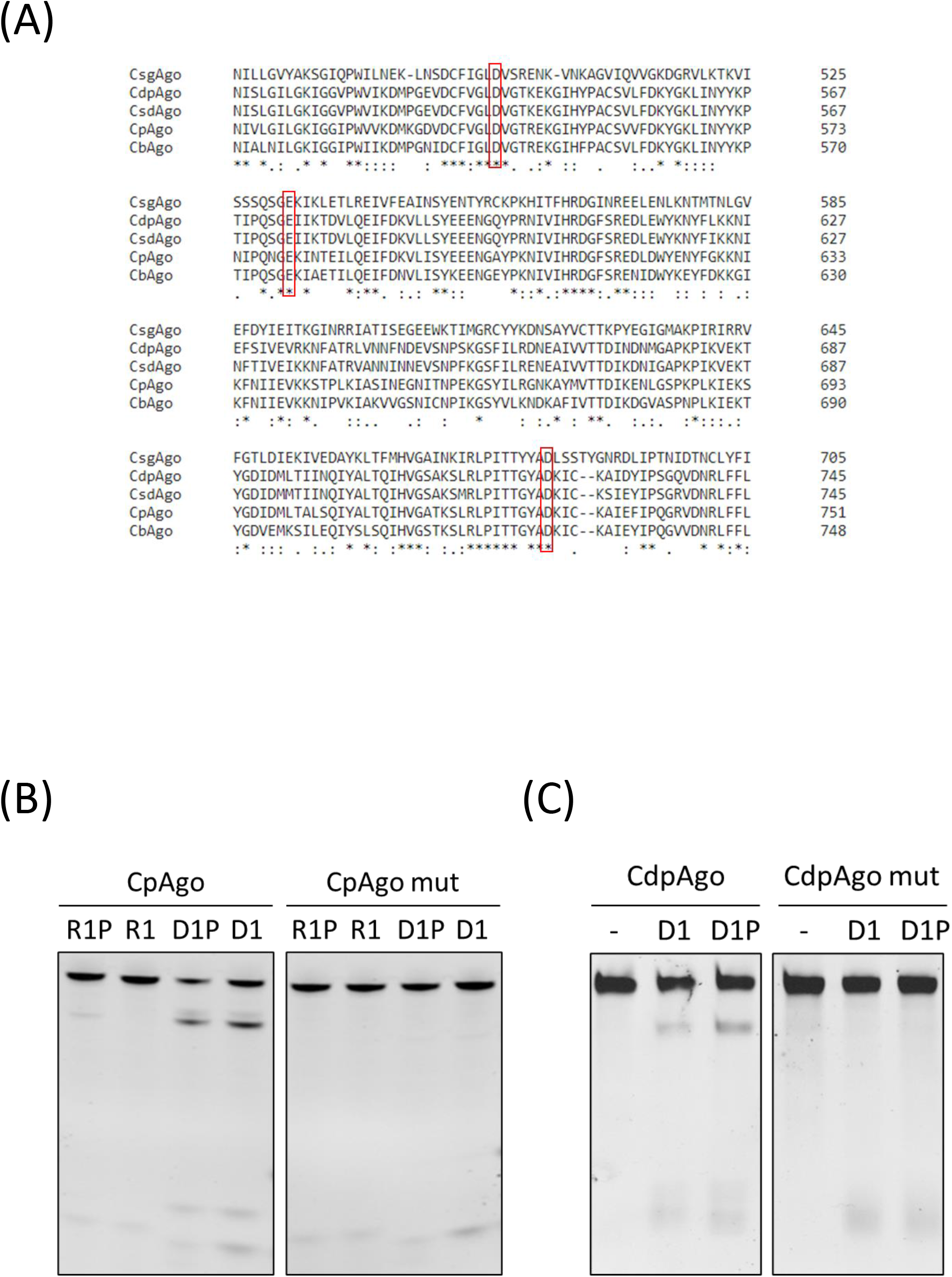
Clostridial pAgos require the integrity of the DEDX motif to maintain nuclease activity. (A) PIWI domains from five Clostridial pAgos were aligned. Catalytic residues (DED) were highlighted in red. (B). Wild-type CpAgo or mutant CpAgo (D544A, E580A and D730A) were expressed and purified as described. Purified pAgos were combined with phosphorylated guide RNA (R1P), unphosphorylated guide RNA (R1), phosphorylated guide DNA (D1P) or unphosphorylated guide DNA (D1) as indicated in the presence of a suitable single-stranded DNA template. (C) Wild-type CdpAgo or mutant CdpAgo (D538A and D608A) were expressed and purified as described. Purified pAgos were combined with phosphorylated guide RNA (R1P), unphosphorylated guide RNA (R1), phosphorylated guide DNA (D1P) or unphosphorylated guide DNA (D1) as indicated in the presence of a suitable single-stranded DNA template.

**Supplementary Figure 4.**
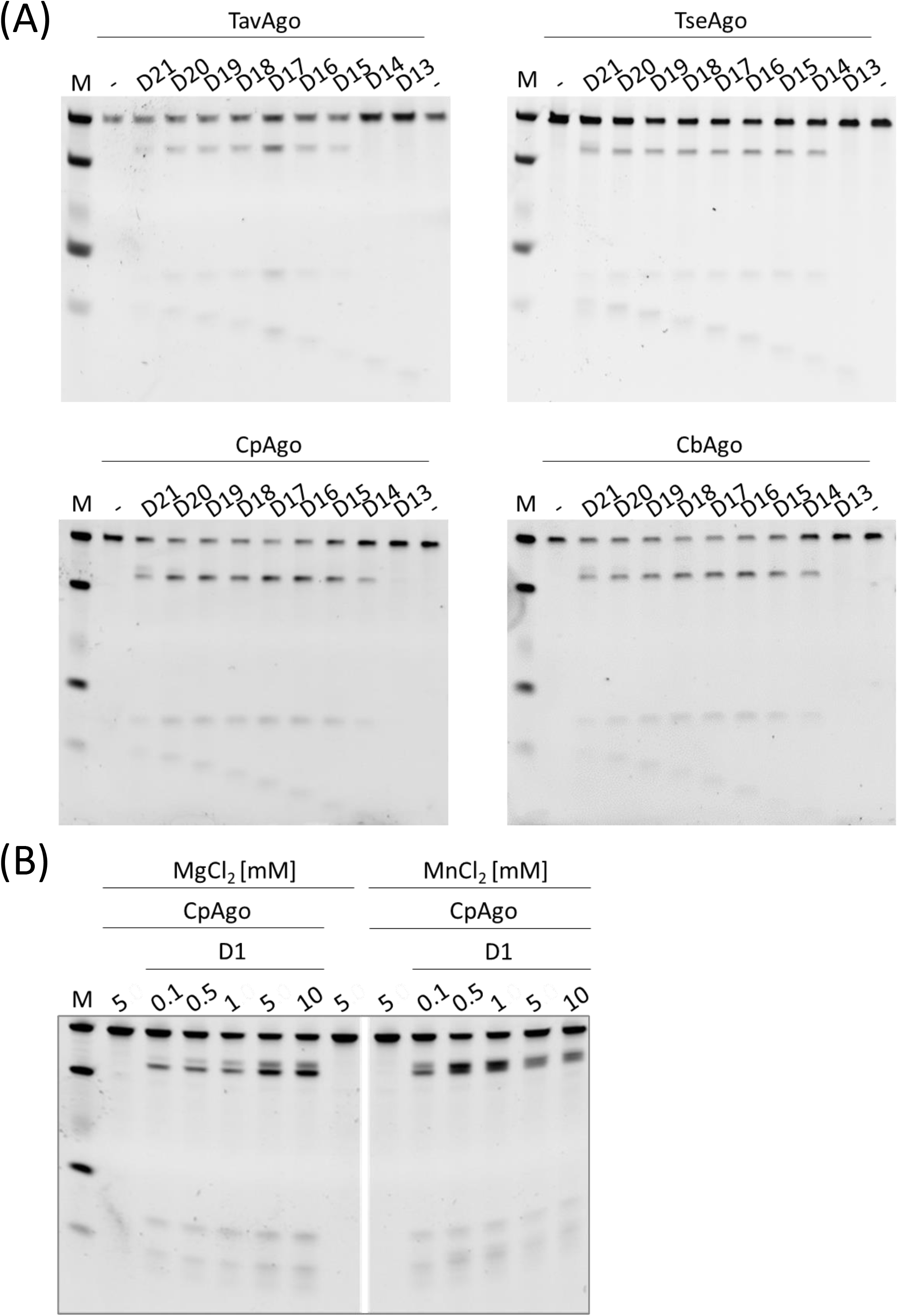
Requirements for nuclease activity of four pAgos. (A) Four pAgos (TavAgo, TseAgo, CpAgo and CbAgo) were obtained as described in Materials & Methods. Purified proteins were combined with phosphorylated guide DNAs of various lengths (see **Supplementary Table 4** for details) and a suitable single-stranded DNA template. Cleavage activity was monitored by TBE urea gel electrophoresis and SYBR Gold staining. (B) Recombinant purified CpAgo was combined with phosphorylated guide DNA D1 and a suitable single-stranded DNA template in the presence of varying concentrations of MgCl2 or MnCl2 (as indicated). Cleavage activity was monitored as described in (A).

**Supplementary Figure 5.**
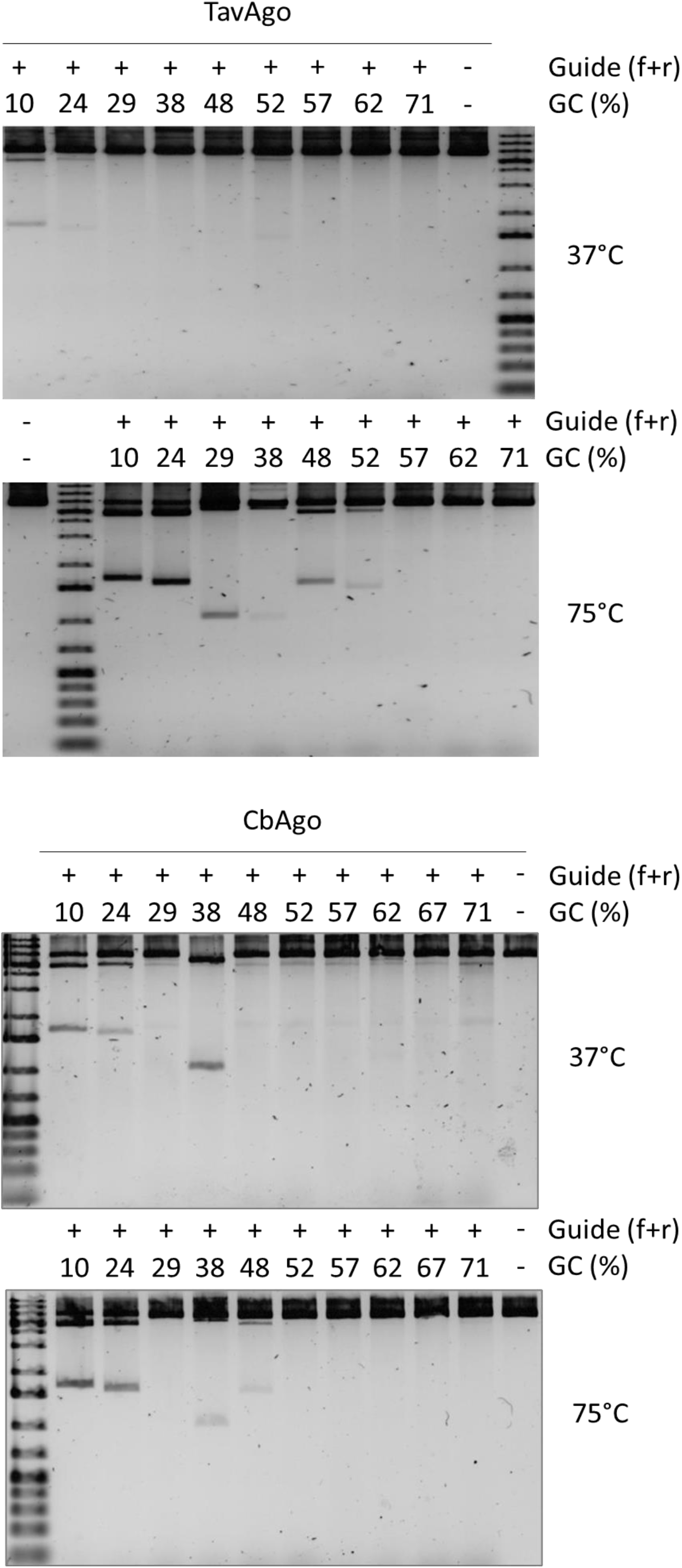
TavAgo and CbAgo-mediated plasmid cleavage in the presence of various guide DNAs. Recombinant purified TavAgo (A) or CbAgo (B) were incubated with plasmid #56 and a set of nine guide DNA pairs harboring increasing GC content (as indicated) in the presence of ET-SSB at 37°C or 75°C (as indicated). Subsequently, plasmids were linearized by addition of a suitable restriction enzyme. Cleavage activity was monitored by agarose gel electrophoresis and SYBR Gold staining.

